## Supplementary material for "Estimating Microbial Population Data from Optical Density": Plate Reader Calibration Protocol

### Preparation

When calibrating a microtiter plate reader for microbial growth experiments, it is essential to begin with a culture in which almost all the particles (cells) can form colonies, i.e., are colony-forming units or CFU. To achieve that, a mid-log phase culture is required - one in which the culture has not begun to enter stationary phase.

There are two easy ways to prepare overnight cultures that are highly reproducible and contain almost exclusively viable cells. Both methods involve limiting cell growth before stationary phase has begun.

#### Method 1: Using rich broth media to obtain oxygen-limited cultures

Inoculate a colony into 10 ml of your favorite broth medium in a 15 ml plug-seal centrifuge tube, then tighten down the plug-seal cap and allow the culture to stand overnight at the preferred growth temperature **without shaking or other method of aeration**.

The details are important. **We do not want the culture to be aerated** because we want to maintain the culture at exponential or early stationary phase. Using the 10 ml of medium in a 15 ml tube allows only 5 ml of oxygen-containing head space. The plug-seal cap ensures that no additional oxygen is available to the growing culture.

#### Method 2: Using nutrient limited mineral salts medium

A mineral salts medium, such as M9 medium, containing a limiting concentration of a carbon source such as glucose (0.02% w/v) works well for many organisms. Inoculate a suitable volume of carbon-limited medium with a single colony and shake well overnight at the optimum growth temperature.

A couple of days before doing the calibration, grow either an oxygen limited or nutrient limited culture. Measure the OD of that culture in 6 wells of a microtiter plate and the OD of the uninoculated medium in another 6 wells (the blanks). Subtract the mean OD of the blank wells from the mean OD of the wells containing the culture. That OD is the *corrected OD* of the nutrient or oxygen limited culture. Use either a 96-well plate or a 384-well plate containing the volume you typically use in that plate. For this purpose, it doesn't matter which size plate you use.

The OD units are OD times volume in ml. For example, a 10 ml culture with a corrected OD of 0.3 contains 3 OD units. The purpose of using OD units is to let you determine the volume of limited culture you will need for the real experiment.

### Calibration

Grow a sufficient volume of limited culture to provide at least 10 OD units. For oxygen limited cultures this may require several 10 ml cultures. If so, combine the cultures. Check the OD of the combined culture to see that there are about 10 OD units.

Spin down the culture(s), pour off the supernatant, and resuspend the pellet in 4.0 ml (or more) of M9 buffer with no nutrients. This volume will depend on how much is needed for the calibration. We found that 4.0 ml was sufficient for both 96-well and 384-well calibrations. Vortex the resuspended cells to mix well. This is the *zero* tube which should have an expected 10 OD units.

### Serial Dilutions for OD Measurements

Prepare a series of tubes numbered 1-10, each containing 2.0 ml of M9 buffer.

Serially transfer 2.0 ml from tube zero to tube 1 and vortex to mix well. Accurate dilutions require both accurate pipetting and thorough mixing at each step. Continue two-fold serial dilutions, transferring and mixing 2 ml at each step.

Fill the wells of a 96 well plate and of a 384 well plate according to the plate layouts below. We use 200 µl per well for 96 well plates and 80 µl per well for 384 well plates, but you can use whatever volumes you choose providing that you use the same volumes for your growth experiments. The layouts provide 6 wells at each dilution

**96 well layout**

| | Wells 1-6 | Wells 7-12 |
|---|---|---|
| **Row A** | Tube zero | Tube 1 |
| **Row B** | Tube 2 | Tube 3 |
| **Row C** | Tube 4 | Tube 5 |
| **Row D** | Tube 6 | Tube 7 |
| **Row E** | Tube 8 | Tube 9 |
| **Row F** | Tube 10 | Buffer blank |
| **Row G** | | |
| **Row H** | | |

If tube zero was at OD = 2.5, then tube 10 will be at about OD = 0.002. In our hands the limit of reliable OD reading is about 0.005, so the serial dilutions will span the useful range of OD.

384 well layout

| | Wells 1-6 | Wells 7-12 | Wells 13-18 | Wells 19-24 |
|---|---|---|---|---|
| **Row A** | Tube zero | Tube 1 | Tube 2 | Tube 3 |
| **Row B** | Tube 4 | Tube 5 | Tube 6 | Tube 7 |
| **Row C** | Tube 8 | Tube 9 | Tube 10 | Buffer blank |

Of course, you can use whatever layout you prefer, but these layouts will allow you to use the accompanying "Calibration Calculator" Excel spreadsheet for your calculations.

#### Read each plate in your microtiter plate reader

Read each plate at 5-minute intervals for 30 minutes, shaking as usual between readings. This allows the plate reader to settle down.

#### Serial dilutions and plating for colony counts

From tube zero, prepare $10^4$-, $10^5$-, $10^6$- and $10^7$-fold dilutions in M9 buffer so that you have at least 1.0 ml of each dilution. Again, pipette carefully and mix well at each dilution. Plate 100µl per plate onto 5 broth plate for each of the $10^4$-, $10^5$-, $10^6$- and $10^7$-fold dilutions. Incubate the plates at the appropriate temperature until colonies have grown.

### Analyze the Data

#### Colony Counts

Count the colonies on whichever dilution has the closest to an average of about 100 colonies per plate. The viable cells per ml in the zero dilution is 10 x dilution x mean colonies per plate.

#### OD Readings

#### 96 well plate, sheet 1

From the plate reader output file copy the 30-minute OD readings into a ***copy*** of the Calibration Calculator excel file, cells D14 through I25.

Into cell A10 enter the dilution that gave closest to 100 colonies per plate. Into cell B10 enter the mean number of colonies per plate for that dilution. Cell C10 will show the cells/ml in the zero tube.

Cells B14 - B24 will show the cells/ml in tubes 0 through 10.

J14-J25 will show the mean ODs in tube 1-10 and the buffer blank.
K14-K24 will show the mean corrected OD, i.e., mean OD - mean OD of the buffer blank.

#### 384 well plate, sheet 2

From the plate reader output file copy the 30-minute OD readings into a ***copy*** of the Calibration calculator excel file, cells D14 through I25.

Into cell A10 enter the dilution that gave closest to 100 colonies per plate. Into cell B10 enter the mean number of colonies per plate for that dilution. Cell C10 will show the cells/ml in the zero tube.

Cells B14 - B24 will show the cells/ml in tubes 0 through 10.

J14-J25 will show the mean ODs in tube 1-10 and the buffer blank.
K14-K24 will show the mean corrected OD, i.e., mean OD - mean OD of the buffer blank.

### Graph Mean Corrected OD vs Cells/ml and Fit Curve

You will need a graphing program that has curve-fitting functions. We use DataGraph for MacOS (https://www.visualdatatools.com/DataGraph/) or KaleidaGraph for both Mac and Windows (https://www.synergy.com/). Other programs include GraphPad Prism (Mac and Windows) (https://www.graphpad.com/scientific-software/prism/) and MagicPlot (Mac, Windows and Unix) (https://magicplot.com/).

Into your graphing program paste the Mean Corrected OD column down through values ≥0.005 and the corresponding Cells/ml. Plot corrected OD (X-axis) vs Cells/ml (Y-axis) and fit a polynomial of degree 4 to those point. This will generate the equation that predicts cells/ml from OD.

The equation will be of the form cells/ml = (A x $OD^4$) + (B x $OD^3$) + (C x $OD^2$) + (D x OD) + E, but will probably be displayed as $AX^4$ _ $BX^3$ + $CX^2$ + DX + E. Some programs display the equation as

E + DX +$CX^2$ + $BX^3$ + $AX^4$. However, it is displayed, in your favorite text editor create a `.cal` file that lists the coefficients, one per line, in the order A, B, C, D, E and be sure that the last line is blank.

We name the file based on the species and the number of wells in the plate; e.g. `Eco384.cal` for E. coli in a 384 well plate.

#### Example to illustrate using the Calibration Calculator

The file `Calibration_384well_Ecoli.xlsx` in the Calibration Tools folder is the output of an *E. coli* calibration of a serial dilution Tube 0 in tubes 1-15, and a buffer blank. The ODs from the 30-minute reading were copied into columns D-I, rows 14-29 of a Calibration Calculator file. The buffer blanks, cells E1-E6, were copied into cells D31-I31. The colony counts from a $10^6$ dilution in which 0.1 ml samples were plated onto 5 LB plates had an average of 266 colonies per plate. The dilution and colony counts were entered in cells A10 and B10 respectively of the Calibration Calculator. The values of the cells/ml were automatically calculated in cells B14-B29. The ODs corrected for the blank were automatically calculated in columns J-O in rows 14-29, and the mean corrected ODs were automatically calculated in column P, rows 14-29.

Experience has shown that ODs < 0.005 are unreliable, so the mean corrected ODs in rows 14-24 were plotted vs the cells/ml in those same rows

The polynomial order 4 fit of cells/ml (Y-axis) vs Mean corrected OD (X-axis) gave the equation cells/ml = 513817 + 1.002e9*x + 1.342e9*x^2 + (-2.301e9)*x^3 + 1.552e9*x^4 with $R^2$ = 1.0

The resulting `.cal` file, `Ecoli384.cal` looks like this:

```
1.552e9
-2.301e9
1.342e9
1.002e9
513817
```
